# Detection of a novel parahenipavirus from northern short-tailed shrews (*Blarina brevicauda* [Say, 1823])

**DOI:** 10.1101/2024.10.03.616414

**Authors:** Sakiho Imai, Mai Kishimoto, Masayuki Horie

**Affiliations:** School of Veterinary Science, College of Life, Environment, and Advanced Science, Osaka Prefecture University, Izumisano, Osaka, Japan; Laboratory of Veterinary Microbiology, Graduate School of Veterinary Science, Osaka Metropolitan University, Izumisano, Osaka, Japan; Osaka International Research Center for Infectious Diseases, Osaka Metropolitan University, Osaka, Japan

## Abstract

Parahenipaviruses (genus *Parahenipavirus*) are non-segmented negative strand RNA viruses belonging to the family *Paramyxoviridae* of the order *Mononegavirales*. Parahenipaviruses have been detected in shrews and rodents, some of which have been reported to be associated with human diseases. Although many parahenipaviruses have been identified thus far, there still exist large phylogenetic gaps in parahenipaviruses, suggesting the existence of yet-to-be-identified parahenipaviruses. In this study, we analyzed public RNA-seq data and identified a novel parahenipavirus from northern short-tailed shrews (*Blarina brevicauda* [Say, 1823]), tentatively named Blarina brevicauda virus (BbV). Based on sequence comparisons between BbV and its most closely related viruses, BbV meets the International Committee on Taxonomy of Viruses species demarcation criteria, and therefore we propose that BbV is a novel species of virus in the genus *Parahenipavirus*. Furthermore, mapping analysis using RNA-seq data derived from multiple tissues of northern short-tailed shrews suggested that BbV is kidney-tropic. This study provides novel insights into the diversity of parahenipaviruses, which would contribute to a deeper understanding of their evolution and control of infectious diseases.

## Introduction

The genus *Parahenipavirus* is a newly established genus within the family *Paramyxoviridae* of the order *Mononegavirales*, consisting of viruses relatively closely related to the members of the genus *Henipavirus* [1]. The genome of parahenipaviruses contains conserved genes typical of paramyxoviruses, including N, P/V/W/C, M, F, G/H/HN, and L, arranged from the 3’ end. Additionally, an additional ORF, X, located between the M and F genes, encodes a putative transmembrane protein that is unique to parahenipaviruses within the family *Paramyxoviridae* [2]. The C, V, and W genes overlap with the P gene; the C protein is derived from an alternative reading frame of P, while the V and W proteins are expressed through mRNA editing, involving the insertion of one or two G residues, respectively.

Parahenipaviruses have been mainly detected in shrews and rodents [2–6], some of which are associated with human diseases. In 2022, Langya virus was detected in febrile patients in China, with shrews identified as the natural reservoir [3]. Similarly, Mojiang virus was detected in rats from an abandoned mine where three patients died of severe pneumonia, although a direct causal link remains unclear [4]. While other parahenipaviruses have not been linked to human disease, their natural hosts, such as rodents and shrews, frequently come into contact with humans, highlighting the need to elucidate the diversity of parahenipaviruses for controlling viral infectious diseases.

The diversity of parahenipaviruses is not fully understood. To date, 12 species and several unclassified viruses have been identified [1]. However, significant phylogenetic gaps within the genus *Parahenipavirus* suggest the existence of many unidentified viruses. Further exploration is needed to deepen our understanding of parahenipavirus diversity and evolution as well as to control infectious diseases.

Sequencing data from diverse samples have been deposited in public databases, some of which may contain virus-derived sequences. Efforts to identify viruses using publicly available data have led to the discovery of many novel viruses, demonstrating the usefulness of re-analyzing archived data for virus discovery [7]. As new public data continue to accumulate over time, they could serve as a valuable resource for identifying additional novel viruses.

In this study, we searched for parahenipaviruses using public RNA-seq data and identified a novel parahenipavirus from the northern short-tailed shrew (*Blarina brevicauda* [Say, 1823]). We determined the nearly complete genome sequence of the virus and tentatively named it Blarina brevicauda virus (BbV). Mapping analyses revealed that viral reads were predominantly detected in kidney-derived data, suggesting that BbV is kidney-tropic. Furthermore, this virus meets the International Committee on Taxonomy of Viruses (ICTV) species demarcation criteria, and we propose BbV as a novel species of virus within the genus *Parahenipavirus*.

## Materials and Methods

### Identification of Blarina brevicauda virus from public RNA-seq data

RNA-seq data (SRR17216496) was downloaded from NCBI SRA [8] and preprocessed using fastp (v0.23.4) [9] with the options “-l 35 -x -y”. The preprocessed reads were then assembled using SPAdes (v4.0.0) [10] with default settings. Contigs of 100 nucleotides or longer were extracted using SeqKit (v2.4.0) [11] and subjected to a BLASTx [12] search against the NCBI clustered nr database [8] on July 29, 2024.

To validate the viral contig, we mapped the RNA-seq data back to the contig using Magic-BLAST (v1.7.2) [13] with the options “-no_unaligned -splice T -word_size 16 -lcase_masking - limit_lookup F”. Read depth at each contig position was calculated using SAMtools (v1.19) [14]. Positions with a read depth of 5 or greater were considered reliable, and regions with a read depth below 5 were removed. The mapping pattern was visualized and manually inspected using Geneious Prime (v2023.2.1, https://www.geneious.com). The final contig, deposited in DDBJ under the accession number BR002473, was used for downstream analyses.

### Annotation of Blarina brevicauda virus genome

For viral gene annotation, ORFs longer than 300 nucleotides were extracted. The putative amino acid sequences of these ORFs were subjected to BLASTp searches against viral sequences (taxid: 10239) in the NCBI nr database, using the options “-word size 2 -evalue 1E-10.” Viral genes were annotated based on the BLAST results.

To identify putative transcription signal sequences, the intergenic regions were extracted and aligned using MAFFT (v7.490) [15] with the E-INS-i algorithm. Based on this alignment and known paramyxovirus signal sequences, the putative transcription signal sequences were manually determined.

The transmembrane domains were predicted using the DeepTMHMM webserver (v1.0.42) [16].

### Phylogenetic analyses

Amino acid sequences of the N and L proteins from BbV, other parahenipaviruses, henipaviruses, and three viruses from other genera within the family *Paramyxoviridae* (Supplementary Table 1) were aligned using MAFFT (v7.490) with the E-INS-i algorithm and ambiguously aligned regions were trimmed with trimAl (v1.4.rev22) [17] using the “-strict” option. Phylogenetic trees were inferred by the maximum likelihood method using RAxML-NG (v1.1.0) [18]. The LG+I+G4+F model, selected by ModelTest-NG (v0.2.0) [19], was used for tree reconstruction. The reliability of the phylogenetic tree was assessed by 1,000 bootstrap resamplings using the transfer bootstrap expectation method. The alignments used for the phylogenetic analyses are available in the Supplementary materials.

### Calculation of amino acid sequence identities for classification

Following ICTV species demarcation criteria (https://ictv.global/ictv/proposals/2023.018M.Paramyxoviridae_reorg.zip [accessed on Sep 22, 2025]), we performed BLASTp searches against viral proteins (taxid: 10239) in the NCBI nonredundant (nr) database using the BbV L-protein sequence as the query, identifying Henipavirus sp. LS-SCDW-1 (PQ421759.1) as a top hit. However, phylogenetic analysis based on L-protein sequences indicated that Ninorex virus (OQ438286.1) is more closely related to BbV. Accordingly, both viral sequences were included in subsequent analyses. Pairwise BLASTp alignments were performed between BbV and Henipavirus sp. LS-SCDW-1 or Ninorex virus for the six major proteins of paramyxoviruses (N, P, M, F, G, and L), and the average amino acid sequence identity was calculated.

### Mapping analysis for detection of Blarina brevicauda virus-derived reads

To detect Blarina brevicauda virus-derived reads, we mapped 306 publicly available RNA-seq datasets obtained from animals in the order *Eulipotyphla* (Supplementary Table 2) to the BbV genome sequence using Magic-BLAST (v1.7.2). The number of mapped reads was then counted using SAMtools (v1.19).

## Results

### Identification of a novel parahenipavirus-like contig

During our extensive search for viruses in public databases, we detected a viral contig from RNA-seq data derived from the kidney of *Blarina brevicauda* (family *Soricidae*, order *Eulipotyphla*). A BLASTx search using this contig revealed that the top hit was the L protein of Ninorex virus (WJL29506.1) [6], a member of the genus *Parahenipavirus*, with 74.6% amino acid sequence identity (Table 1).

**Table 1.** One-on-one BLASTp comparison between Blarina brevicauda virus and Ninorex virus.

| Protein | Identity (%) |  |
| --- | --- | --- |
|  | Ninorex virus | Parahenipavirus sp.<br>LS-SCDW-1 |
|  | OQ438286.1 | PQ421759.1 |
| N | 50.7 | 73.8 |
| P | 34.7 | 44.6 |
| M | 76.2 | 79.4 |
| F | 55.7 | 62.4 |
| G | 33.4 | 48.6 |
| L | 74.6 | 75.8 |
| Average | 54.2 | 64.1 |

To evaluate the reliability of the contig, we mapped the RNA-seq data from which the contig was derived to the viral contig sequence. Positions with a read depth of 5 or more were considered reliable, and sequences that did not meet this threshold were removed. The final contig length was 16,645 nucleotides. We named this virus Blarina brevicauda virus (BbV).

### Annotation of the Blarina brevicauda virus genome

Next, we determined the genome organization of BbV, which contains nine ORFs. Using the amino acid sequences of these ORFs as queries, we performed BLASTp analysis and found sequence similarity to the N, P/V/W/C, M, ORF X, F, G, and L genes of other parahenipaviruses (Fig. 1a).

**Figure 1.**
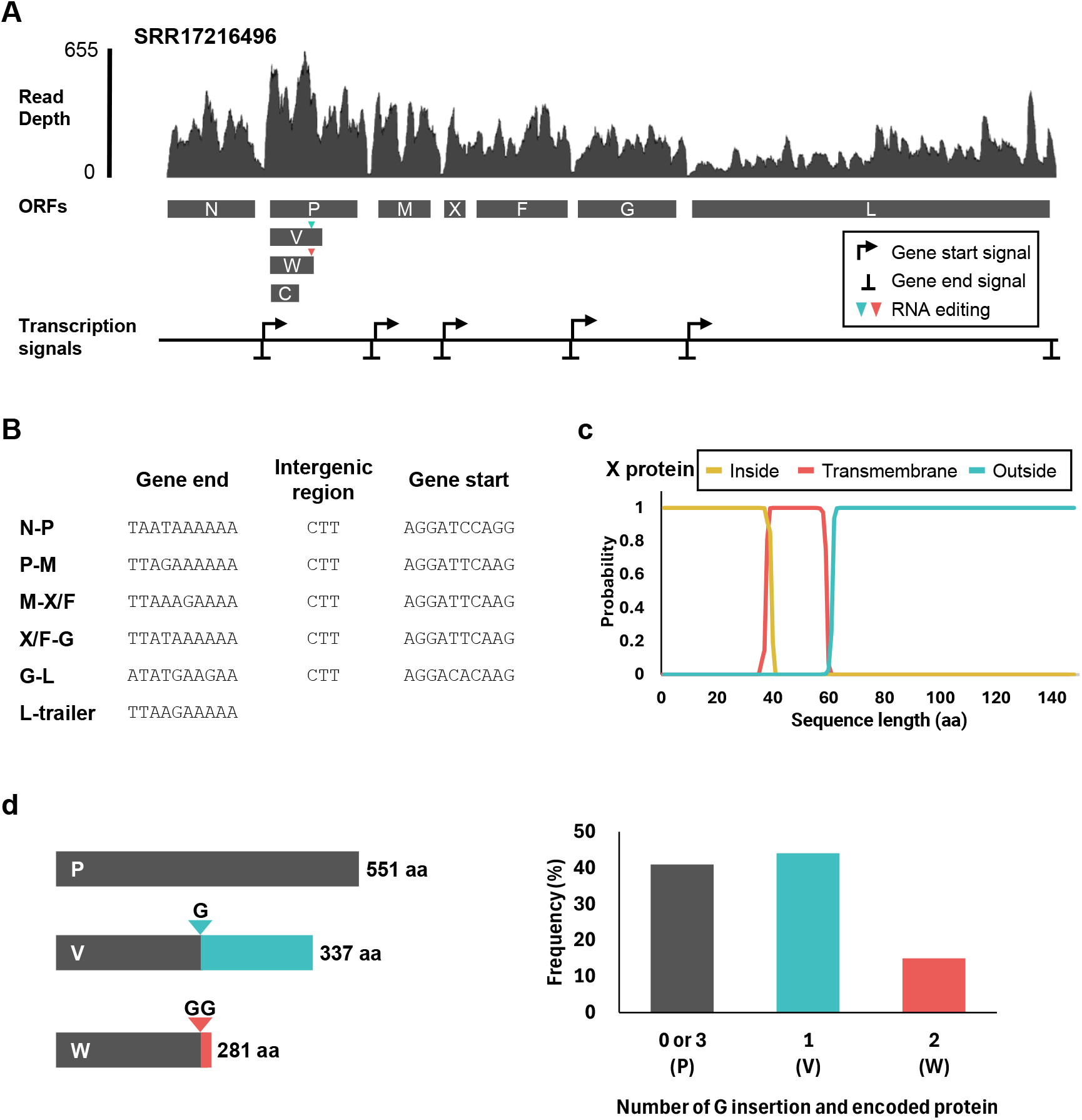
Genome organization of Blarina brevicauda virus (BbV). **(A)** Schematic representation of the open reading frames (ORFs) and transcription signal sequences of BbV. **(B)** Predicted transcription signal sequences of BbV. **(C)** Prediction of the transmembrane domain of the ORF X protein using DeepTMHMM. **(D)** Schematic diagram illustrating the P, V, and W proteins (left) and the average frequencies of guanine base insertions (right). Detailed data are provided in Supplementary Table 3.

To identify the transcription signal sequences, we aligned the intergenic regions and manually determined the sequences, referencing those of other orthoparamyxoviruses (Fig. 1b). The intergenic CTT motif, conserved in most orthoparamyxoviruses, was also present in all BbV intergenic regions. The putative transcription start and end signals were identified as AGGAYHCARG and WTADRRARAA, respectively. Notably, the mapping data showed a marked decrease in read depth in the intergenic regions (Figs. 1A and S1), suggesting active viral transcription in the samples.

ORF X, located between the M and F genes, is a distinctive gene of parahenipaviruses and is predicted to encode a transmembrane protein. Using DeepTMHMM, we predicted the transmembrane region of BbV’s ORF X to be between amino acids 39-60, which is similar to other parahenipaviruses (Fig. 1C).

Paramyxoviruses typically utilize mRNA editing through the insertion of G nucleotide(s) by transcriptional slippage to express P, V, and W proteins. To investigate whether BbV employs the same mechanism, we reanalyzed the BbV mapping data. We detected G nucleotide insertions at position 2,720 on the BbV genome (Figs. 1A and 1D), which result in the production of three different proteins: the P protein (551 aa) without insertion, the V protein (337 aa) with a single G insertion, and the W protein (281 aa) with two G insertions. Based on the mapping data, the frequencies of expression for these proteins were 40.9%, 44.1%, and 15%, for P, V, and W, respectively (Fig. 1D and Supplementary Table 3).

### Evolutionary analyses on Blarina brevicauda virus

To elucidate the evolutionary relationship between BbV and other parahenipaviruses, we performed phylogenetic analyses using the N and L proteins (Fig. 2A and B). In both phylogenetic trees, BbV clustered with Ninorex virus, Wufeng Chodsigoa smithii henipavirus 1, and a recently detected parahenipavirus LS-SCDW-1, all of which were detected in animals from the family *Soricidae* (Fig. 3).

**Figure 2.**
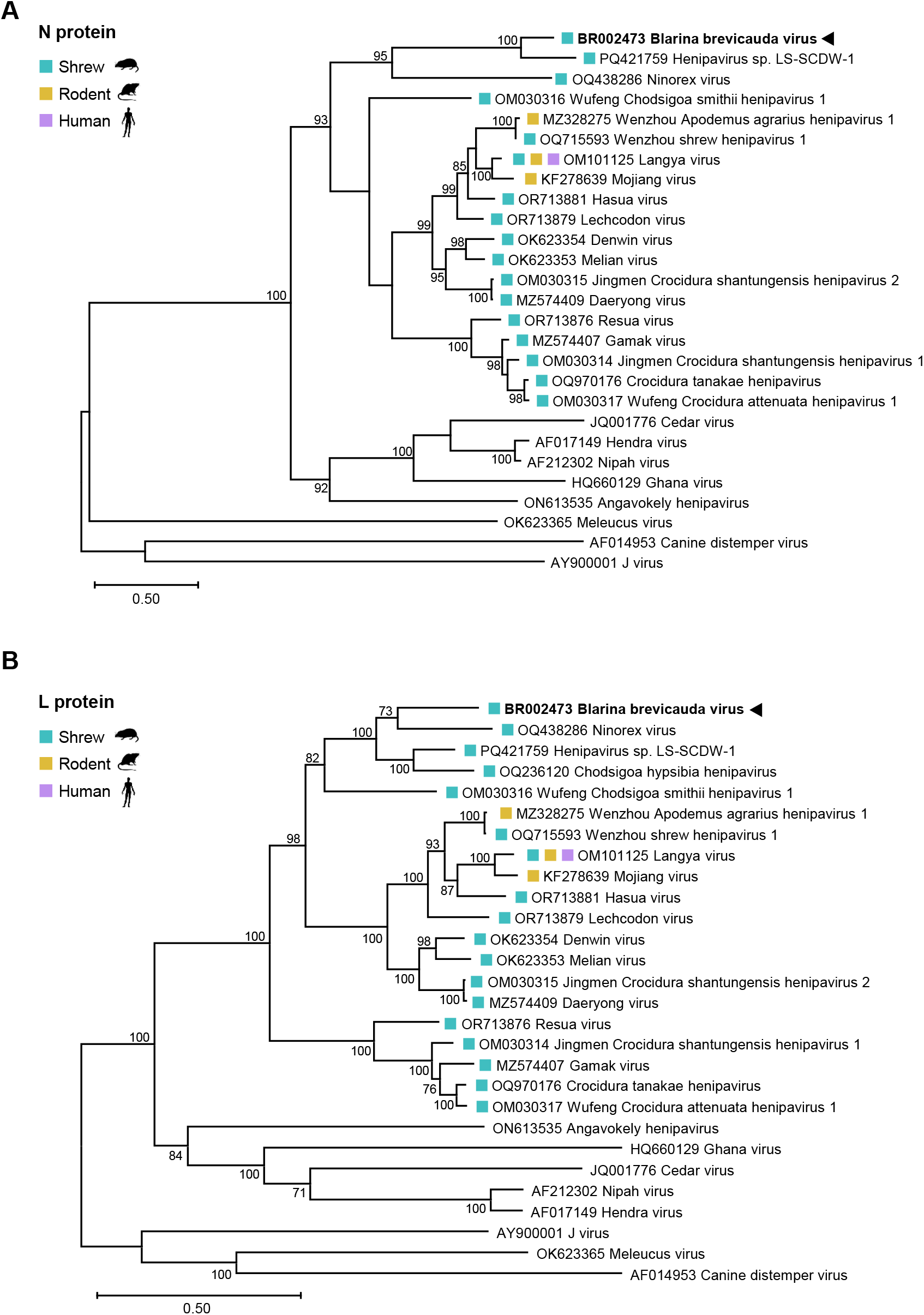
Phylogenetic trees of parahenipaviruses, henipaviruses, and other paramyxoviruses. Maximum likelihood phylogenetic trees were inferred using amino acid sequences of the N **(A)** and L **(B)** proteins. Bootstrap values below 70 are not shown. The scale bar represents the number of amino acid substitutions per site.

**Figure 3.**
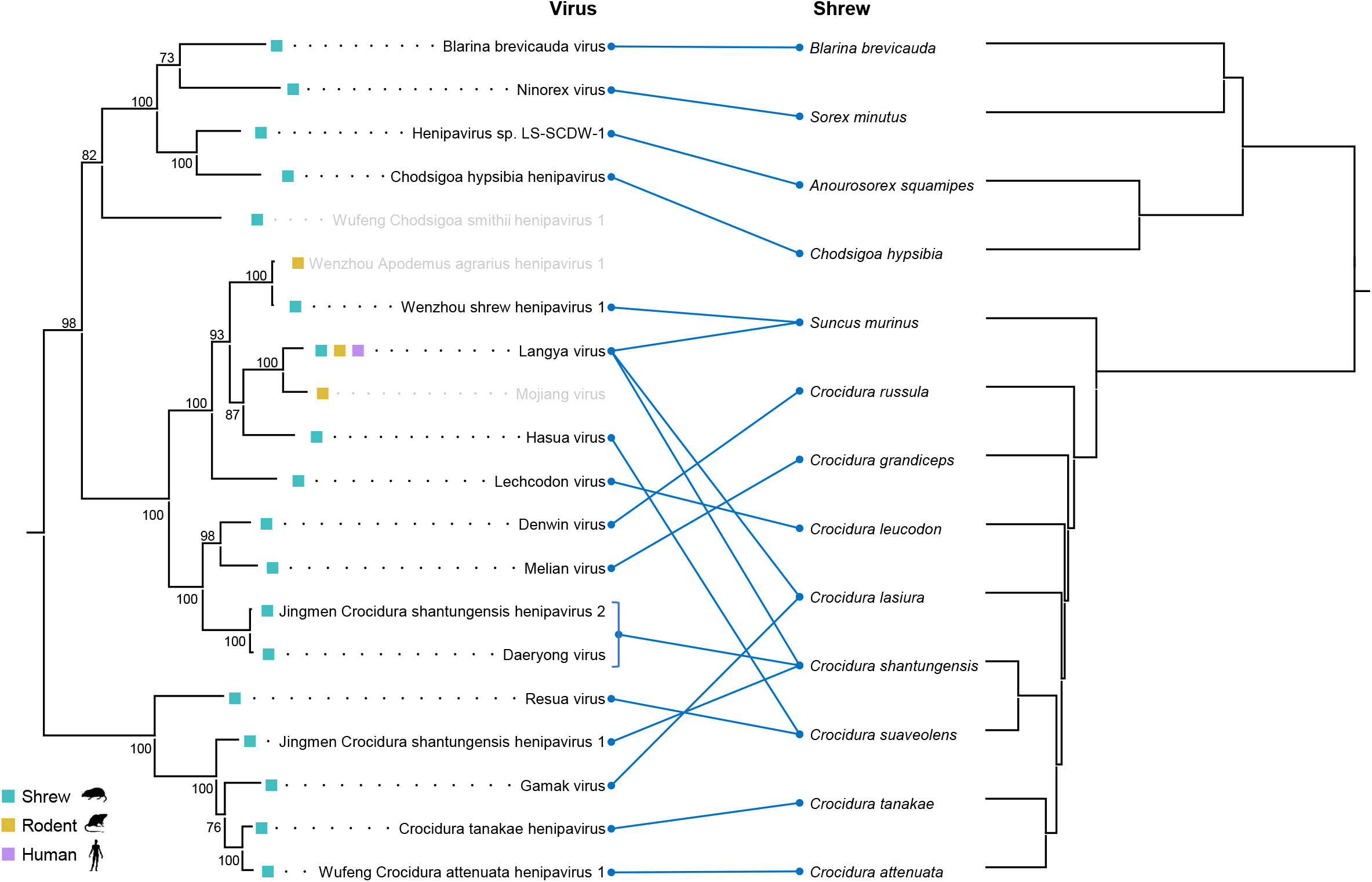
Comparison of topologies between viral and host phylogenies. Phylogenetic trees of viruses (left) and hosts (right) are shown; the viral tree is adapted from Fig. 2B, and the host tree was obtained from TimeTree 5 [28]. Blue lines link each virus to the host species. Viruses lacking a corresponding host lineage in TimeTree 5 or detected in non-shrew hosts are shown in gray.

We then calculated amino acid sequence identities between BbV and its closest relatives, Ninorex virus and Parahenipavirus sp. LS-SCDW-1, for the six major paramyxovirus proteins (N, P, M, G, F, and L). The sequence identity for each protein is shown in Table 1; on average, BbV shares 54.2% identity with Ninorex virus and 64.1% with Parahenipavirus sp. LS-SCDW-1.

### Parahenipaviruses may have largely evolved in co-divergence manner with the hosts

To further understand the evolution of parahenipaviruses, we compared the topologies of trees between parahenipaviruses and their hosts. Our analysis showed that the topology of the viral phylogeny is largely congruent with that of the host phylogeny, although some incongruences exist (Fig. 3) (see details in Discussion).

### Comprehensive detection of BbV reads in public RNA-seq data

To further characterize BbV, we conducted a comprehensive search for BbV-derived reads in public RNA-seq data through mapping analysis. We mapped 306 RNA-seq datasets from animals of the order *Eulipotyphla*, which includes *Blarina brevicauda*, the putative host of BbV, to the viral contig and counted the mapped reads. As a result, BbV-derived reads were detected in four additional RNA-seq datasets (Table 2), all of which belong to the same BioProject as the one where BbV was initially identified. Among the five RNA-seq datasets positive for BbV (including the initial dataset), four were derived from the kidney and one from the skin. The number of viral reads in the kidney-derived datasets was markedly higher than in the skin-derived dataset. Notably, while this BioProject also contains data from the brain, heart, liver, and skin, no BbV-derived reads were detected in brain, heart, or liver; only one skin dataset contained BbV reads (Table 2).

**Table 2.** Sample information and mapped read numbers in BioProject PRJNA788430.

| **SRA Run accession** | **Sample name** | **Mapped read number** | **Read per million** | **Tissue** |
| --- | --- | --- | --- | --- |
| SRR17216479 | Shrew217 | 0 | 0 | Brain |
| SRR17216480 |  | 0 | 0 | Heart |
| SRR17216481 |  | 14,304 | 124.2 | Kidney |
| SRR17216482 |  | 0 | 0 | Liver |
| SRR17216483 |  | 0 | 0 | Skin |
| SRR17216484 | Shrew218 | 0 | 0 | Brain |
| SRR17216485 |  | 0 | 0 | Heart |
| SRR17216486 |  | 7,081 | 51.89 | Kidney |
| SRR17216487 |  | 0 | 0 | Liver |
| SRR17216488 |  | 0 | 0 | Skin |
| SRR17216489 | Shrew220 | 0 | 0 | Brain |
| SRR17216490 |  | 0 | 0 | Heart |
| SRR17216491 |  | 18,978 | 170.74 | Kidney |
| SRR17216492 |  | 0 | 0 | Liver |
| SRR17216493 |  | 35 | 0.26 | Skin |
| SRR17216494 | Shrew221 | 0 | 0 | Brain |
| SRR17216495 |  | 0 | 0 | Heart |
| SRR17216496 |  | 24,085 | 186.08 | Kidney |
| SRR17216497 |  | 0 | 0 | Liver |
| SRR17216498 |  | 0 | 0 | Skin |

## Discussion

Currently, 12 species and several unassigned viruses are reported in the genus *Parahenipavirus* [1], but large phylogenetic gaps remain, and the diversity of parahenipaviruses has yet to be elucidated. In this study, we identified the nearly complete genome sequence of a novel paramyxovirus, BbV, from RNA-seq data of *Blarina brevicauda*. Notably, our analysis showed that the average amino acid sequence identities of the six major viral proteins (N, P, M, F, G, and L) between BbV and its closest relatives, Ninorex virus and parahenipavirus LS-SCDW-1, are 54.2% and 64.1%, respectively. This meets the ICTV species demarcation criteria for the family *Paramyxoviridae* (average identity below 85%; https://ictv.global/ictv/proposals/2023.018M.Paramyxoviridae_reorg.zip [accessed on Sep 22, 2025]). Based on these findings, we propose that BbV is a novel species of virus within the genus *Parahenipavirus*.

Our data strongly suggest that *B. brevicauda* is the authentic host of BbV. In metaviromic analyses, viral contamination from various sources can sometimes occur, complicating the identification of authentic hosts. However, our mapping analysis identified a significant number of virus-derived reads in the kidney. Further, we showed a marked decrease in read depth in the intergenic regions, suggesting that the virus actively transcribed in the samples. Additionally, closely related parahenipaviruses have also been detected in shrews [2,3,5,6,20,21]. These findings indicate that the virus is actively transcribed in the kidney, strongly suggesting that *B. brevicauda* is the authentic host of BbV.

Parahenipaviruses may have largely evolved in co-divergence manner with the hosts. Our analysis indicates that the topology of the viral tree is congruent with that of the host one, at least at the genus level (Fig. 3). For viruses detected from *Crocidura* shrews, some incongruences were observed, but this could be attributed to the difficulty of resolving relationships within *Crocidura* because the diversification of those shrews seems to have occurred over a very short timescale [22]. Therefore, our data suggest that parahenipaviruses have largely co-diverged with their hosts. On the other hand, several parahenipaviruses have been detected in multiple host species, indicating cross-species transmission. In particular, some viruses within the lineage represented by Langya virus have been detected in species other than shrews (including rodents and humans), suggesting cross-species transmission from shrews to more distantly related animals. Note that evidence of cross-species transmission of parahenipaviruses has so far been found in viruses in a limited lineage. Further identification of parahenipaviruses filling the phylogenetic gap would provide insights into the evolution of parahenipaviruses.

Our data also suggest that BbV is kidney-tropic. A total of 20 RNA-seq datasets from *B. brevicauda* are registered under BioProject PRJNA788430, from which BbV was detected. These datasets are derived from five different tissues (brain, skin, heart, kidney, and liver) apparently from four individuals according to the metadata. Our mapping analysis revealed that viral reads were almost exclusively found in the kidneys of all four individuals. Although viral reads were detected in the skin of one individual, the quantity was significantly lower than in the kidney. Thus, the viral reads in the skin sample might be due to contamination and/or index hopping. Additionally, Hasua virus, Resua virus, Lechcodon virus, and Denwin virus, parahenipaviruses detected from shrews, were suggested to be kidney-tropic [23]. Furthermore, many parahenipaviruses were detected in the kidneys [2,5,6,24]. Thus, many of the members of the genus *Parahenipavirus* may be kidney-tropic.

This study is the first to report the frequency of G-base insertions through RNA editing of the P gene in parahenipaviruses. The transcripts encoding P, V, W proteins accounted for 40.9%, 44.1%, and 15% of the total on average, respectively (Fig. 1D). However, the ratio of P/V/W transcripts is known to be changed depending on the infection stage [25,26]. Further analyses of infected animals and infection experiments will provide more detailed insights into the RNA-editing of parahenipaviruses.

During the preparation of this manuscript, an independent study reporting similar findings was published [20]. However, that study classified the virus within the genus *Henipavirus*, and therefore several taxonomic and related descriptions were inaccurate, as noted by Haring et al. [27]. Importantly, another paper also classified a parahenipavirus (isolate LS-SCDW-1; PQ421759.1) as a member of the genus *Henipavirus* [21]. In the present work, we analyze the virus as a member of the genus *Parahenipavirus*, providing a more accurate understanding of this virus.

In this study, we identified a novel paraphenipavirus from RNA-seq data derived from *B. brevicauda*, contributing to a deeper understanding of the diversity of parahenipavirus. However, significant phylogenetic gaps remain, and the characteristics of BbV are not yet fully understood. Further epidemiological and virological studies are necessary to better understand this virus.

## Acknowledgments

The super-computing resources were provided by Human Genome Center, the Institute of Medical Science, the University of Tokyo and the NIG supercomputer at ROIS National Institute of Genetics.

This study was supported by KAKENHI grant numbers JP21H01199 (MH), JP22K19234 (MH), JP23K20902 (MH), JP24K21922(MH), and JP24K18455 (MK) and the 2024/2025 Osaka Metropolitan University (OMU) Strategic Research Promotion Project (Young Researcher) (MK).

## Notes

### Competing Interest Statement

The authors have declared no competing interest.

### Summary of Updates

Figure 1 revised, Figure 2 revised, Figure 3 added; Table 2 revised, Supplementary files updates.

